# Quantitative evaluation of warming and hemodilution as two techniques to increase the flow rate of blood during simulated massive transfusion of packed red blood cells

**DOI:** 10.1101/2022.01.29.478293

**Authors:** Netra Shah

## Abstract

Massive bleeding is a leading cause of deaths in trauma, surgeries, wartime injuries, and childbirth. In these situations of ongoing blood loss, it is critical for patients to be rapidly resuscitated. Currently, patients who require major transfusions are given cold and concentrated blood from the blood bank. This blood does not flow fast due to its viscosity. Equipment for rapid resuscitation such as rapid transfusers are very costly and are not accessible everywhere. Additionally, specialized training is required to operate these machines. Therefore, it is necessary to determine simple, quick, and effective ways to increase the flow rate of blood. The flow rate can be increased by warming and hemodilution. The purpose of this study was to quantify the effects of these simple and quick methods since they can be implemented anywhere with easily accessible equipment. In this experiment, expired packed red blood cells were warmed and diluted. The time the blood took to flow through the tubing was measured and a flow rate was calculated for each trial. The research hypothesis stated that diluted blood at 39°C would have the fastest flow rate. The research hypothesis was supported and there was a 366% in flow rate by warming and diluting blood. There was a 232% increase by simply diluting the blood and a 170% increase by warming. Six t-tests were conducted for this data and they were all significant at a level of 0.001. These results indicate the significant effects of warming and diluting blood and this research could improve patient care during emergencies and massive blood loss.

## Introduction

Currently, one of the leading causes of death is trauma and 40% of trauma deaths are caused by bleeding [1]. Other situations where patients need to receive blood rapidly are when they have major blood loss from surgery, dehydration, or shock [2]. Massive blood loss requires rapid resuscitation. The high morbidity and mortality rates in trauma could be reduced by increasing the flow rate of blood during transfusion. Equipment, such as rapid transfusers, are not available everywhere and are very expensive. The average cost of this equipment is $10,000-20,000 and the tubing and supplies for each use are very expensive. In addition, people who use these machines need to have specialized training. Furthermore, IV access is often limited in these emergencies. Hence, there is a need to implement quick and easy techniques to increase flow rate of blood that can be implemented in most situations with easily available equipment.

Two simple, yet effective, ways are hemodilution and warming. These methods can increase flow rate substantially and should be possible to implement anywhere, thereby saving many lives. The purpose of this study was to model the impact of these techniques using expired human packed red blood cells and commonly available IV tubing and cannulas which simulate real-life clinical scenarios.

Although medicine has had major advances, the effects of hemodilution and warming have not been studied and quantified as a means to increase flow rate of blood. Previous studies have evaluated in vitro methods to increase the flow rate of IV fluids by increasing cannula size, increasing height of the IV tubing system, and the use of pressure bags [3]. There have been studies determining the effect of different intravenous access devices on flow rate of IV fluid [4]. Other areas of research include determining the safe limits of hemodilution to maintain blood gas exchange and the effect of hematocrit on the flow rate of blood in different organs, such as the brain and kidneys. In a study performed by Hudak, Koehler, Rosenberg, Traystman, and Jones, it was found that cerebral blood flow decreases as hematocrit increases [5]. This study quantifies the effect of these two simple and quick methods in a simulated real-life experience.

Blood has many important functions in the body, such as carrying oxygen and nutrients to all the cells in the human body. For massive transfusions, patients are given packed blood from the blood bank. The blood banks maintain blood in a cold and concentrated form. Hence, the flow rate of the blood will be very slow. It was hypothesized that increasing the blood flow rate would be a better and more efficient way to rapidly resuscitate.

The independent variables in the study were hematocrit and temperature of the blood. Poiseuille’s law states that the flow of any liquid is inversely proportional to its viscosity [6]. Hence to enhance flow rate, the viscosity of blood must be reduced. The two main ways to achieve this are reducing the hematocrit or increasing the temperature. The hematocrit of blood stored in the blood bank in the form of packed red blood cells (PRBCs) is about 60-70%. In contrast, the hematocrit of normal blood is about 35-45%. According to Klabunde, “When the hematocrit is 40%, the relative viscosity of blood is 4 times that of water and when the hematocrit is 60%, the relative viscosity is 8 times more than the viscosity of water” [6]. By diluting the blood (hemodilution), the viscosity of the blood reduces, which improves circulatory hemodynamics. According to Messmer and Kemming, healthy volunteers may tolerate a hematocrit of around 20% in absence of disease [7].

The viscosity of blood could be also reduced by increasing the temperature of the blood. The temperature of stored blood is about 2 – 6°C, while the normal body temperature is 37°C. If the blood is warmed up to normal body temperature, then the flow rate will increase. Cold blood can also be harmful to patients due to high lactate and potassium levels. Normal blood has a pH of 7.40, while cold blood is more acidic [8]. Giving cold blood to patients during a transfusion could also cause hypothermia and acid-base abnormalities.

The dependent variable in this study was the flow rate of the blood that is being transfused. For this specific experiment, the flow rate measures how fast the blood will flow through IV tubing and catheters to resemble a massive transfusion. The flow rate of blood in this experiment was measured in mL/min.

The four levels of the independent variable are packed red blood cells (referred as blood in rest of article) at 4°C, diluted blood at 4°C, blood at 39°C, and diluted blood at 39°C. Blood at 4°C was chosen as the control. A higher temperature of 39°C was chosen because normal human body temperature is 37°C and the maximum temperature that blood warmers can warm blood is 42°C [9]. Blood that is 39°C and diluted and blood that is 4°C and diluted are the other two IV levels. The blood was diluted, by adding normal saline solution, from a hematocrit of 60% to 40%. The hematocrit was changed to 40% because in normal adults, the normal hematocrit is 38% - 50% [10].

## Materials and methods

### Preparation of specimen

The experiment was studied in a BSL-2 Lab and protective eyewear, gloves and scrubs were worn to ensure safety. Since no human subjects were involved and no human specimens were collected for this study, approval from an ethics committee and participant consent were not needed. Three units of expired packed red blood cells from the blood bank mixed together in a one-liter bag. This blood was divided equally into four transfusion bags to ensure each bag had the same amount of blood with the same hematocrit that was found to be 60. The four bags were labeled “Blood at 4°C”, “Blood at 39°C”, “Blood at 39°C and diluted”, and “Blood at 4°C and diluted” with a sharpie and the bags at 4°C were stored on ice. The hematocrit of the blood was measured, and it was 60. The amount of normal saline that was needed to reduce the hematocrit to 40 was calculated using 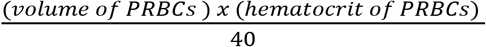. Thus, 100 mL of saline was added to 200 mL of blood in the two bags labeled diluted and the final hematocrit was measured and was 40.

### Attachment of tubing and setup

A standard blood collection bag was attached to standard blood transfusion tubing and the bag was labeled “Bag A”. An 18-gauge IV catheter was attached to the other side of the tubing. The blood collection bag was measured and made sure that it was 120 centimeters above the IV-catheter. The other end of the IV catheter was attached to another blood collection bag. This bag was labeled “Bag B”.

## Experiment

### A. Measuring flow rate using blood at 4 degrees celsius

Fifty mL of blood at 4°C was aliquoted using a 50 mL syringe from the bag labeled “Blood at 4°C.” A volume of fifty mL was chosen for each trial to ensure that the temperature did not change during the experiment. The blood was placed into Bag A. The roller clamp of Bag A was opened. The time it took to empty Bag A was measured in min:sec using a phone timer. The flow rate was calculated and recorded using the formula mL/time taken (min). Using a 50 mL syringe, 50 mL of blood was removed from Bag B and placed into Bag A. Bag B was kept in ice to maintain the temperature at 4°C. The blood was reused for the entire experiment. The entire procedure was repeated 24 more times.

### B. Measuring flow rate using blood at 39 degrees Celsius

The blood from the bag labeled “Blood at 39°C” was warmed to 39°C using a blood warmer set at 39°C. Bag B was placed in a water bath that maintained a temperature of 39°C. The entire procedure from “Measuring Flow Rate using Blood at 4°C” was repeated 25 times using the bag labeled “Blood at 39 degrees Celsius” and the results were recorded.

### C. Measuring flow rate using diluted blood at 39 degrees Celsius

The procedure done in “Measuring Flow Rate using Blood at 39°C” was repeated 25 times using the bag labeled “Blood at 39°C and diluted” and the results for each trial was recorded.

### D. Measuring flow rate using diluted blood at 4 degrees Celsius

The bag labeled “Blood at 4°C and diluted” was stored in an icebox to remain at 4°C. The procedure done in “Measuring Flow Rate using Blood at 4 degrees Celsius” was repeated 25 times using the bag labeled “Blood at 4 degrees and diluted” and the results for each trial was recorded.

## Results

The effects of the temperature and hematocrit of the blood were tested on the flow rate of blood. For each level of the independent variable, a mean was determined. The results of the statistical analysis are shown in Table 3. Figure 1 shows the results in pictorial form. The blood at 39°C and diluted had the highest mean of 46.0 mL/min, while blood at 4°C had the lowest mean (12.6 mL/min). Blood at 4°C and diluted had the second highest mean of 29.2 mL/min and blood at 39°C had the third highest of 21.4 mL/min.

**Figure 1.**
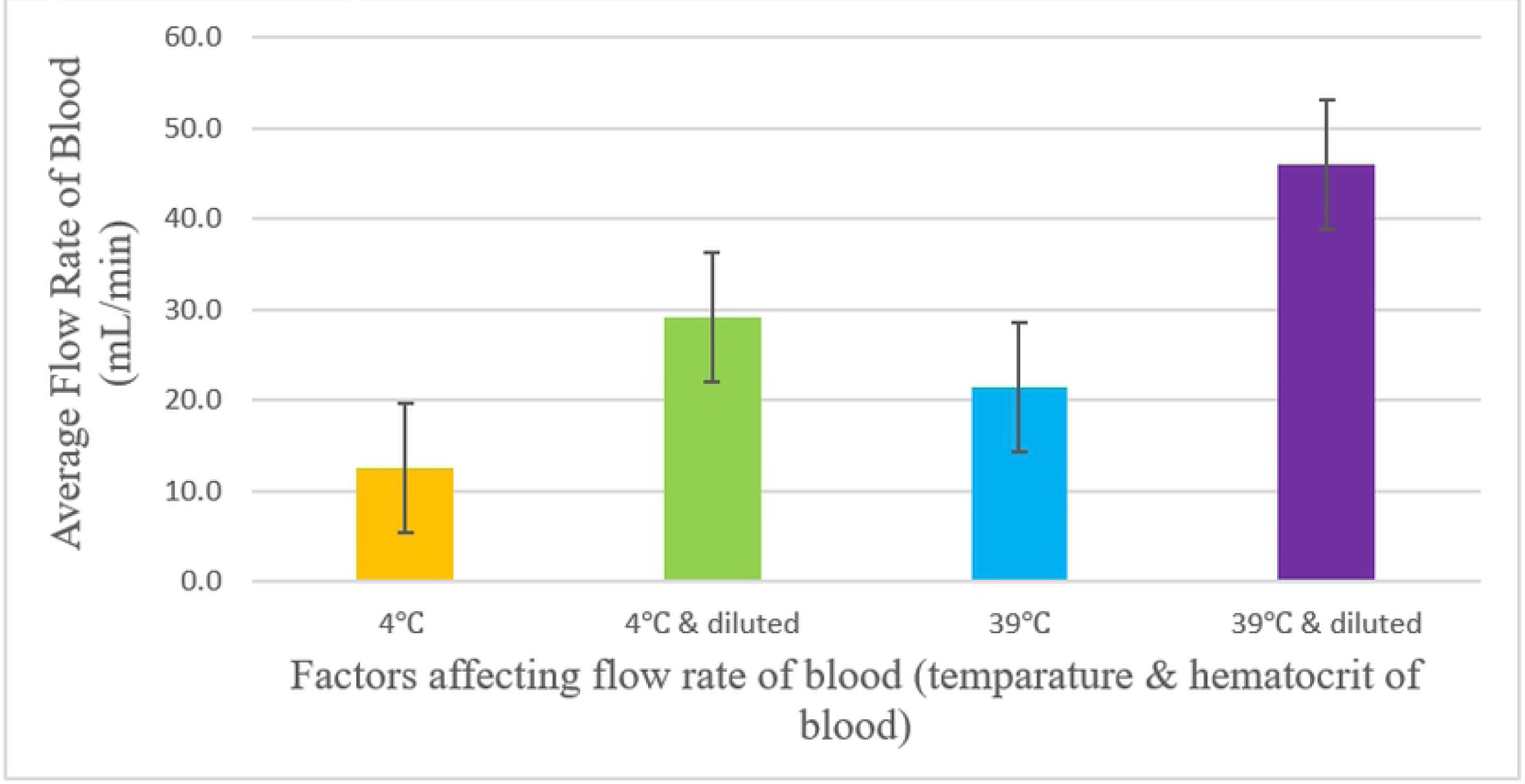
Visual representation of mean flow rate for each IV level.

**Figure 2.**
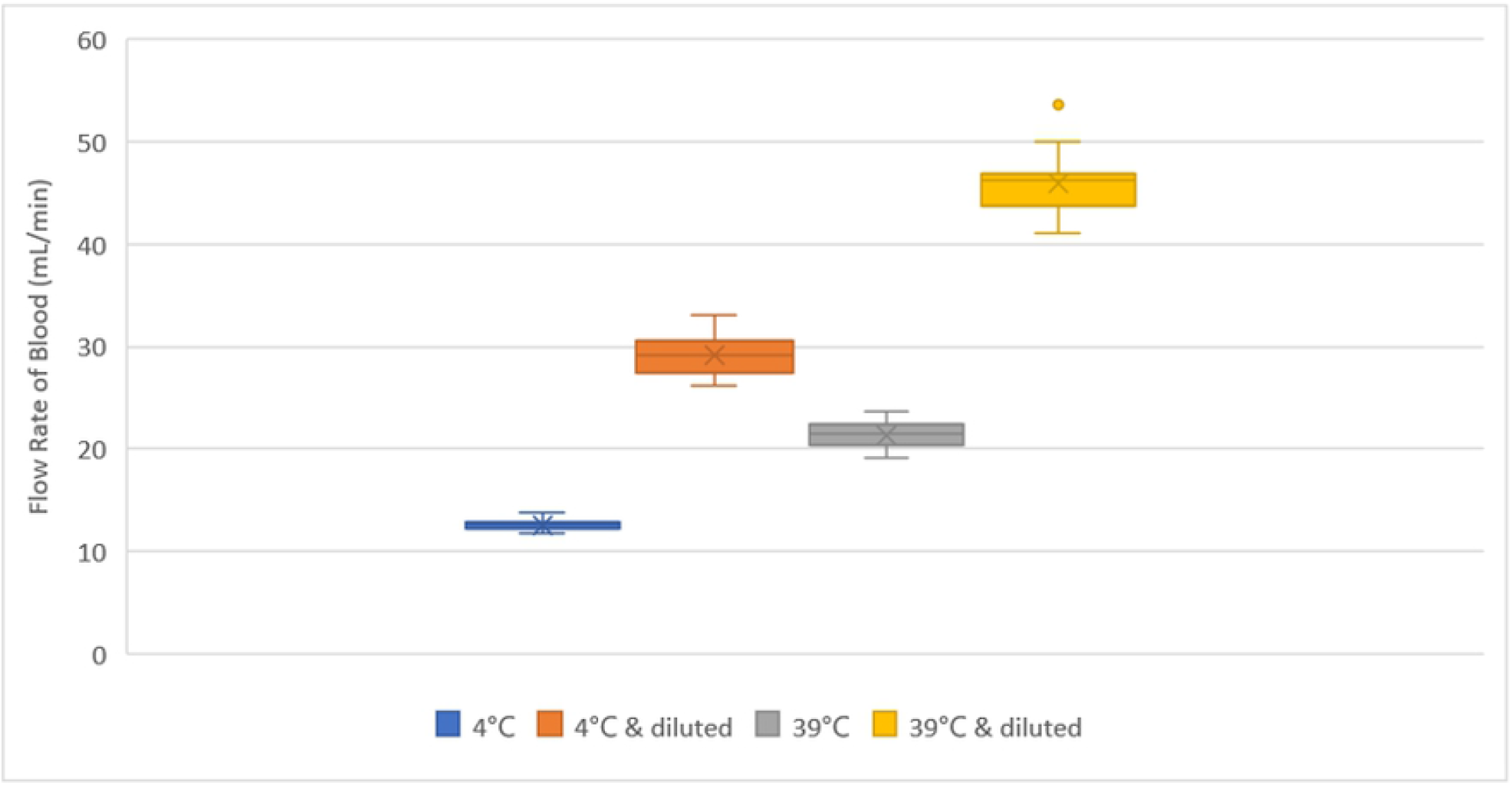
Box and whisker plot on flow rate of blood of four IV levels.

**Table 1.** Time measured for 50 mL blood to flow through the IV tubing.

| Trials | Flow rate of Blood (min:sec) |  |  |  |
| --- | --- | --- | --- | --- |
|  | 4°C | 4°C & diluted | 39°C | 39°C & diluted |
| 1 | 3:54 | 1:55 | 2:33 | 1:09 |
| 2 | 3:53 | 1:50 | 2:16 | 1:04 |
| 3 | 3:52 | 1:55 | 2:09 | 1:00 |
| 4 | 3:59 | 1:54 | 2:14 | 1:02 |
| 5 | 3:55 | 1:43 | 2:16 | 1:04 |
| 6 | 4:06 | 1:49 | 2:07 | 1:04 |
| 7 | 4:13 | 1:54 | 2:20 | 1:04 |
| 8 | 3:53 | 1:38 | 2:08 | 1:13 |
| 9 | 3:39 | 1:44 | 2:26 | 1:05 |
| 10 | 4:11 | 1:43 | 2:23 | 1:05 |
| 11 | 4:00 | 1:38 | 2:22 | 1:04 |
| 12 | 3:43 | 1:49 | 2:23 | 1:04 |
| 13 | 4:07 | 1:36 | 2:30 | 1:05 |
| 14 | 3:55 | 1:38 | 2:24 | 1:06 |
| 15 | 3:38 | 1:45 | 2:28 | 1:05 |
| 16 | 4:00 | 1:45 | 2:30 | 1:12 |
| 17 | 3:54 | 1:40 | 2:28 | 1:08 |
| 18 | 3:58 | 1:43 | 2:36 | 1:09 |
| 19 | 4:02 | 1:34 | 2:37 | 1:07 |
| 20 | 4:04 | 1:38 | 2:17 | 1:04 |
| 21 | 4:05 | 1:42 | 2:19 | 1:10 |
| 22 | 4:07 | 1:31 | 2:14 | 0:56 |
| 23 | 4:13 | 1:33 | 2:18 | 1:10 |
| 24 | 3:57 | 1:41 | 2:10 | 1:02 |
| 25 | 4:14 | 1:44 | 2:13 | 1:04 |
| Mean | 3:58 | 1:43 | 2:20 | 1:05 |

**Table 2.** Calculated flow rate for each IV level.

| Trials | Flow rate of Blood (mL/min) |  |  |  |
| --- | --- | --- | --- | --- |
|  | 4°C | 4°C & diluted | 39°C | 39°C & diluted |
| 1 | 12.8 | 26.1 | 19.6 | 43.5 |
| 2 | 12.9 | 27.3 | 22.1 | 46.9 |
| 3 | 12.9 | 26.1 | 23.3 | 50.0 |
| 4 | 12.6 | 26.3 | 22.4 | 48.4 |
| 5 | 12.8 | 29.1 | 22.1 | 46.9 |
| 6 | 12.2 | 27.5 | 23.6 | 46.9 |
| 7 | 11.9 | 26.3 | 21.4 | 46.9 |
| 8 | 12.9 | 30.6 | 23.4 | 41.1 |
| 9 | 13.7 | 28.8 | 20.5 | 46.2 |
| 10 | 12.0 | 29.1 | 21.0 | 46.2 |
| 11 | 12.5 | 30.6 | 21.1 | 46.9 |
| 12 | 13.5 | 27.5 | 21.0 | 46.9 |
| 13 | 12.1 | 31.3 | 20.0 | 46.2 |
| 14 | 12.8 | 30.6 | 20.8 | 45.5 |
| 15 | 13.8 | 28.6 | 20.3 | 46.2 |
| 16 | 12.5 | 28.6 | 20.0 | 41.7 |
| 17 | 12.8 | 30.0 | 20.3 | 44.1 |
| 18 | 12.6 | 29.1 | 19.2 | 43.5 |
| 19 | 12.4 | 31.9 | 19.1 | 44.8 |
| 20 | 12.3 | 30.6 | 21.9 | 46.9 |
| 21 | 12.2 | 29.4 | 21.6 | 42.9 |
| 22 | 12.1 | 33.0 | 22.4 | 53.6 |
| 23 | 11.9 | 32.3 | 21.7 | 42.9 |
| 24 | 12.7 | 29.7 | 23.1 | 48.4 |
| 25 | 11.8 | 28.8 | 22.6 | 46.9 |
| Mean | 12.6 | 29.2 | 21.4 | 46.0 |

**Table 3.** Statistical analysis with t-tests values.

| Descriptive Information | Factors (temperature & dilution) (mL/min) |  |  |  |
| --- | --- | --- | --- | --- |
|  | 4°C | 4°C and diluted | 39°C | 39°C and diluted |
| Mean | 12.6 mL/min | 29.2 mL/min | 21.4 mL/min | 46.0 mL/min |
| Range | 2.0 mL/min | 6.9 mL/min | 4.5 mL/min | 12.5 mL/min |
| Maximum | 13.8 mL/min | 33.0 mL/min | 23.6 mL/min | 53.6 mL/min |
| Minimum | 11.8 mL/min | 26.1 mL/min | 19.1 mL/min | 41.1 mL/min |
| Variance | 0.279 | 3.768 | 1.709 | 7.149 |
| Standard Deviation | 0.528 | 1.941 | 1.307 | 2.674 |
| 1 SD | 12.072-13.128 | 27.259 - 31.141 | 20.093-22.707 | 43.326-48.674 |
| 2 SD | 11.544-13.656 | 25.318 - 33.082 | 18.786-24.014 | 40.652-51.348 |
| 3 SD | 11.016-14.184 | 23.377 - 35.023 | 17.479-25.321 | 37.978-54.022 |
| Number | 25 | 25 | 25 | 25 |
| Results of a t-test | 4 degrees C vs 4 degrees C and diluted $t = 41.258$ $p < 0.001$<br>4 degrees C vs 39 degrees C $t = 31.206$ $p < 0.001$<br>4 degrees vs 39 degrees and diluted $t = 61.275$ $p < 0.001$<br>4 degrees C and diluted vs 39 degrees $t = 21.640$ $p < 0.001$<br>4 degrees C and diluted vs 39 degrees C and diluted $t = 25.423$ $p < 0.001$<br>39 degrees C vs 39 degrees C and diluted $t = 41.327$ $p < 0.001$<br>At $df=48$; $\alpha = 0.001$; $t = 3.551$ for significance | | | |

The above results indicate that hemodilution had a more significant effect on the flow rate of blood than warming in this experiment. This was suggested because the diluted blood at 4°C had a larger mean than blood at 39°C.

Blood at 39°C and diluted had the highest standard deviation of 2.674, and blood at 4°C had the lowest standard deviation of 0.258. All standard deviations were relatively low. There were two outliers for the control, blood at 4°C, (13.7 and 13.8) and there was one outlier for blood at 39°C and diluted (53.6). Since both approaches led to an increase in flow rate, the research hypothesis was supported. Six t-tests were performed for this experiment at a significance level of 0.001. The calculated values from all six t-tests were larger than the table t value (3.551). These results imply that the null hypothesis should be rejected and that these two approaches cause an increase in flow rate of expired PRBCs.

## Discussion

The purpose of the experiment was to investigate quick, yet simple maneuvers to substantially increase the flow rate of blood during a massive, rapid transfusion. To simulate real life situations, commonly accessible equipment in hospitals were used. The flow rate of blood is substantially dependent on the blood’s viscosity, which can be manipulated by temperature and hematocrit changes. Normal saline solution was used to dilute the blood.

This experiment showed that the mean flow rate at 4°C was 12.6 ml/min and at 39°C, it was 21.4 ml/min. This showed a 170% increase by simply warming the blood. The flow rate of 4°C and diluted blood had a 232% increase, showing the impact of hemodilution. When the blood was warmed and diluted, it had a flow rate change of 366%. Multiple t-tests were done for this experiment and all the t-tests determined that the results were statistically significant at a level of significance of 0.001. The degrees of freedom for all the t-tests was 48. The results of this experiment imply that all the data was due to the independent variable, and not chance. Since the results were due to the independent variable, the null hypothesis was rejected.

Many researchers around the world have studied the effect of blood temperature on the viscosity of blood. According to Cinar, Senyol, and Duman, increasing the temperature of blood from 36.5°C to 39.5°C, decreased the viscosity by about 10%, increasing the flow rate of blood [11]. Other studies have shown when the hematocrit was increased by about 20%, it caused 56% of the decrease in the flow rate of blood in the brain [5]. These studies have all shown that if the blood temperature was increased or if the hematocrit was decreased, the flow rate of blood was increased. The findings in the experiment were similar to the findings in these experiments above since when the temperature was increased or when the hematocrit was decreased, the flow rate of blood increased.

The blood from the blood bank is very concentrated and has a high viscosity. Hence, it does not flow fast through IV tubing. To remedy this, the blood is diluted with 0.9% normal saline. The normal saline acts like a volume expander and the blood becomes less viscous [12]. In this experiment, the blood was hemodiluted, so it flowed faster through the tubing. Another factor which affects the viscosity is the temperature of blood. As the temperature is lowered, the blood becomes thicker and hence it flows slowly. According to Klabunde, “viscosity increases about 2% for each degree centigrade decrease in temperature” [6]. The packed red blood cells that are given to the patients are stored at about 4°C. When the blood is warmed up to 40°C from 4°C, there is a 36-degree change which corresponds to a 72% reduction in the viscosity. The findings in this experiment were significant since the flow rate increase 170% with warming and 232% with hemodilution.

Although promising, this approach is not without challenges. Residual blood in the tubing is possible and could affect repeating trials. Small blood clots may also form because the IV tubing is a non-natural material. Challenges also arise from maintaining precise temperature control and starting times during the experiment. An automated machine may solve these problems. Finally, the impact of the size of IV catheters, the type of IV tubing, and the location of the IV catheter on the flow rate of blood should be investigated.

## Conclusion

Bleeding is a very common cause of morbidity and mortality in the world. Massive transfusions are often needed in these situations. Specialized equipment for massive transfusions is very limited and expensive. Warming and hemodilution are two techniques that dramatically increase the flow rate of blood. These methods are very simple, safe, quick, require commonly accessible equipment and can be implemented in most places. Warming and hemodilution could potentially save many patients’ lives.

## Acknowledgements

I would like to thank Dr. Shamaskin MD and Umesh R. Desai, Ph.D. for their guidance throughout the process of conducting the experiment and proofreading the paper.

